# Deep sequencing analysis of the circadian transcriptome of the jewel wasp *Nasonia vitripennis*

**DOI:** 10.1101/048736

**Authors:** Nathaniel J. Davies, Eran Tauber

## Abstract

The study of the circadian clock has benefited greatly from using *Drosophila* as a model system. Yet, accumulating evidence suggests that the fly might not be the canonical insect model. Here, we have analysed the circadian transcriptome of the Jewl wasp *Nasonia vitripennis* by using RNA-seq in both constant darkness (DD) and constant light (LL, the wasps are rhythmic in LL with period shortening). At a relatively stringent FDR (q < 0.1), we identified 1,057 cycling transcripts in DD and 929 in LL (fraction of 6.7% and 5.9% of all transcripts analysed in DD and LL respectively). Although there was little similarity between cycling genes in *Drosophila* and *Nasonia*, the functions fulfilled by cycling transcripts were similar in both species. Of the known *Drosophila* core clock genes, only *pdp1e*, *shaggy* and *Clok* showed a significant cycling in *Nasonia*, underscoring the importance of studying the clock in non-model organisms.

## Introduction

The circadian clock regulates fundamental biological processes such as sleep (Huang *et al*. 2011), metabolism (Huang *et al*. 2011), and the immune system (Scheiermann *et al*. 2013), and has implications for a wide range of human diseases. Notable examples of diseases linked to the circadian clock include cancer (Kelleher *et al*. 2014), Alzheimer’s disease (Musiek *et al*. 2015), cardiovascular disease (Takeda and Maemura, 2011), obesity (Maury *et al*. 2010), diabetes (Maury *et al*. 2010), and depression (Quera Salva, *et al*. 2011). A primary output of the clock is circadian regulation of transcription, a trait which has been demonstrated in mammals (Hughes *et al*. 2009), insects (McDonald and Rosbash, 2001a), plants (Schaffer *et al*. 2001), and even bacteria (Woelfle and Johnson, 2006). Therefore, analysing transcriptional oscillations in clock controlled genes (CCGs) is a key step in understanding how the daily rhythms produced by the clock are ultimately linked to behavioural phenotypes.

The genetic mechanisms underlying the animal circadian clock were first elucidated through studies of model animals; primarily the fruit fly *Drosophila*. The first clock gene to be identified, period (per), was discovered through mapping the genetic basis of *Drosophila* mutants with aberrant locomotor and eclosion rhythms (Konopka and Benzer, 1971). The discovery of period was followed by the discovery of its heterodimeric partner timeless (tim) (Sehgal *et al*. 1994). These two genes are joined by a roster of other genes working together to produce robust internal rhythms.

The discoveries made in *Drosophila* have been instrumental for understanding the mechanisms of the circadian clock in mammals (Yu and Hardin, 2006). As the principal insect model, *Drosophila* has been used to great effect to model circadian phenomena in humans (Rosato *et al*. 2006). However, as circadian research into non-drosophilid insects has advanced, several alternative clock models have been proposed (Yuan *et al*. 2007), some of which may better model aspects of the mammalian clock than *Drosophila*.

For example, a major difference between the various clock models in insects concerns the light input pathway. The main light input to the clock in *Drosophila* is mediated through cryptochrome (cry1) which is activated in response to light (Ceriani *et al*. 1999), binds to and promotes the degradation of tim (Busza *et al*. 2004), ultimately resulting in the degradation of per (Ko *et al*. 2002, Grima *et al*. 2002). In contrast, mammalian-like cryptochrome (cry2) is not light-sensitive (Yuan *et al*. 2007), but is a part of the core transcriptional feedback loop suppressing its own transcription (and that of per) by interfering with the actions of the CLK-BMAL1 heterodimer (Kume *et al*. 1999, Jin *et al*. 1999). Mammals also lack a homolog for timeless, possessing only a homolog of the *Drosophila* gene timeout (Benna *et al*. 2000), a gene whose potential role in the clock is less clear and less crucial than that of timeless (Gustafson and Partch, 2015, Benna *et al*. 2010).

The Lepidoptera harbour both types of cryptochrome (*Drosophila*-like cry1 and mammal-like cry2) (Tomioka and Matsumoto, 2010), as well as homologs of timeless and timeout (Tomioka and Matsumoto, 2015). The two cryptochromes have been shown to act in a similar way to their *Drosophila* and mammal counterparts; cry1 functions as a light receptor and cry2 serves as a transcriptional repressor (Zhu *et al*. 2008).

Of the major insect orders, the Hymenoptera arguably possess the most mammalian-like core clock architecture, possessing cry2 and timeout but neither cry1 nor timeless (Tomioka and Matsumoto, 2015, Yuan *et al*. 2007). In addition to these molecular similarities, there is evidence that the transcriptional profiles of these genes match more closely the mammalian model than the *Drosophila* model (Rubin *et al*. 2006). Light-entrained circadian rhythms have been demonstrated in the Hymenoptera, but the question of light detection in the Hymenopteran clock remains an open one.

*Nasonia* vitripennis is a parasitoid wasp, which as a research model offers advantages over other hymenopterans, including a fully sequenced genome (Werren *et al*. 2010), systemic RNAi (Lynch and Desplan, 2006), a robust and well-characterised circadian response (Bertossa *et al*. 2013), a fully functional DNA methylation kit (Park *et al*. 2011), and a history as a model for photoperiodism (Saunders, 1969).

In this study, we advance *Nasonia* as an alternative circadian model by using RNA-seq to profile whole-transcriptome gene expression in the *Nasonia* head. As the *Nasonia* clock free-runs in both constant darkness and constant light (Figure 1), we profiled both of these conditions to examine how the two circadian transcriptomes differ. To our knowledge, this is the first circadian RNA-seq study performed in an insect other than *Drosophila*, and the first study to profile the circadian transcriptome oscillating under constant light.

**Figure 1.**
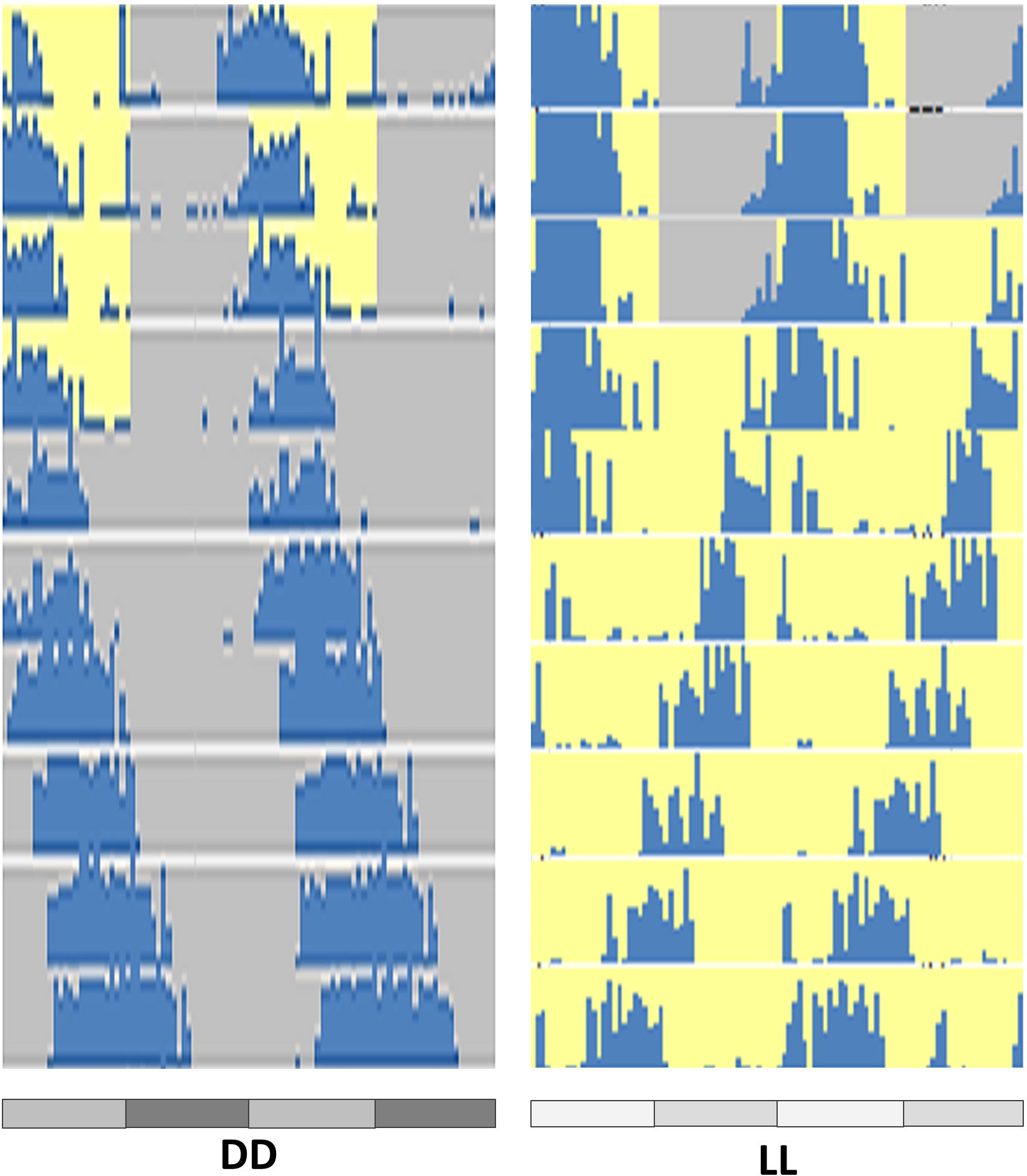
Free-run behavioural rhythm in *Nasonia*. Representative actograms of individual *Nasonia* males in DD (left) and LL (right). Activity counts were sorted into 30 minute bins and plotted in blue. Yellow and grey backgrounds indicate lights on and lights off respectively. Gray and black bars below the actogram indicate the 12 hr subjective day and night.

## Results

### Identifying rhythmic transcription

We first performed an unbiased clustering analysis to ascertain the kinds of expression patterns present in the data. To this end, Mfuzz (Kumar and E Futschik, 2007) was used to carry soft c-means clustering, a method which is less sensitive to biological noise than traditional clustering (Futschik and Carlisle, 2005). After filtering (see Methods), thirty clusters were generated for each condition (Supplementary figures S1 and S2), revealing a variety of potentially rhythmic and non-rhythmic expression trends. Potential asymmetric wave forms were detected in LL (e.g. Supplementary figure S2, clusters 22 and 26).

To identify rhythmic transcripts, we used the RAIN algorithm (Thaben and Westermark, 2014). At false discovery rate (FDR) threshold of 0.1 we identified 1,057 rhythmic transcripts in DD and 929 in LL (Table S1, S2).

Rhythmic transcripts (q < 0.1) were sorted by phase, peak shape, and significance, and plotted (Figure 2A). Examining the phase distribution (Figure 2B), it is apparent that the majority of transcripts show peak expression early in the subjective morning/afternoon or in the subjective night, with fewer transcripts peaking at intermediate times. This disparity in phase is greater in the transcripts which show rhythmic expression in both DD and LL; less than 12% of transcripts in DD and less than 5% in LL show peak expression at intermediate times (Figure 2B). The majority of these transcripts (~87%) exhibit a similar (+−4 hrs) phase in LL to their phase in DD.

**Figure 2.**
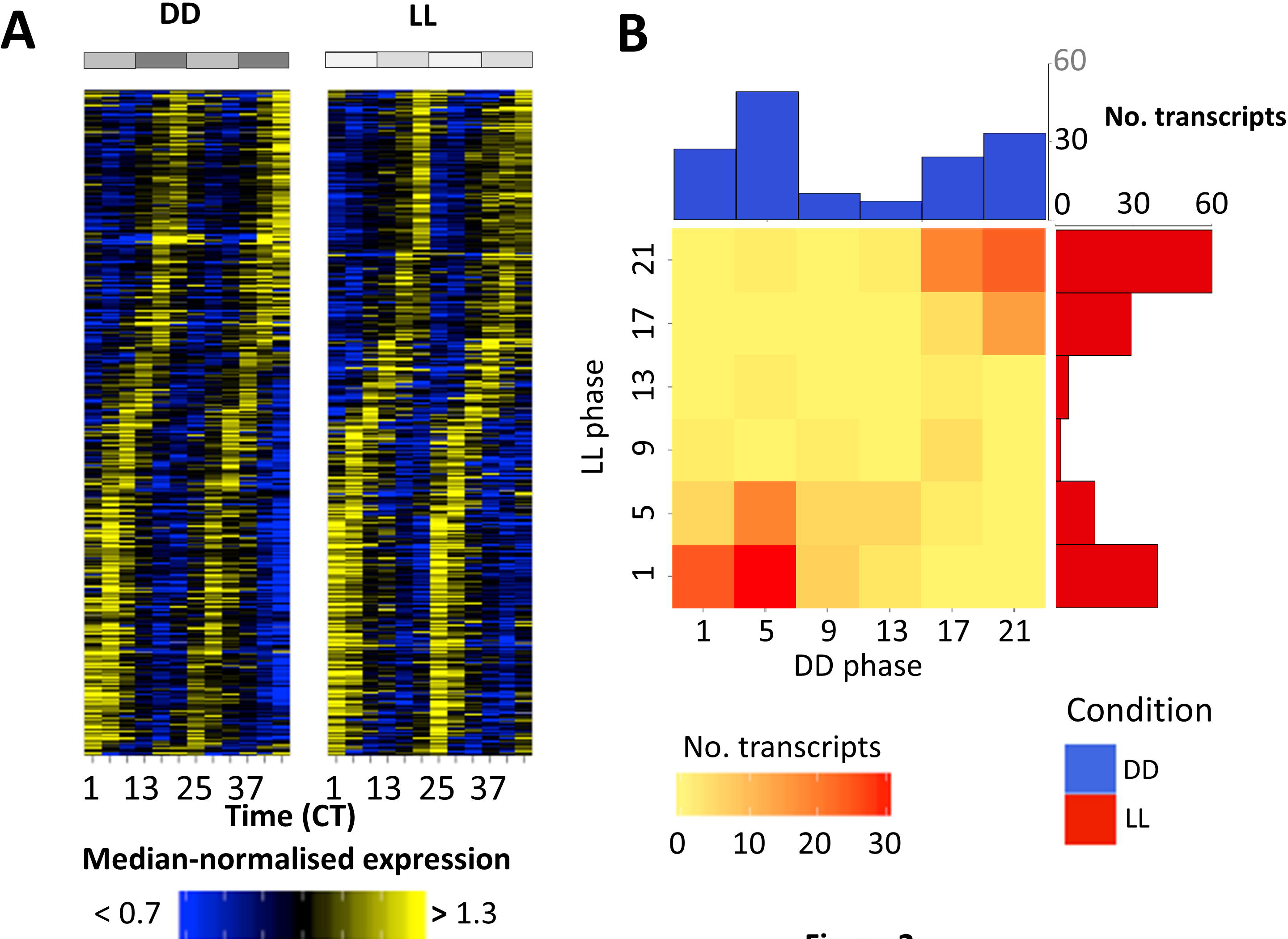
Circadian transcriptional rhythms. (**A**) Heatmap of median-normalised expression of rhythmic (q < 0.1) transcripts in both constant darkness and constant light. (**B**) Histograms and heatmap of phases of rhythmic transcripts (q < 0.1 in both conditions), showing bimodal phase distribution and overlap between the two conditions.

Similarly to *Drosophila* (Hughes *et al*. 2012) and mammals (Hughes *et al*. 2009), the majority of transcripts show only small cyclic changes in expression amplitude over the day; over 80% of reliably quantified (see Methods) transcripts in both conditions have amplitudes (peak expression divided by trough expression) of 2 or less. In both DD and LL, transcripts with exceptionally high amplitudes (> 4) are transcripts with unusually low or high measurements at isolated time-points with no obvious specific shared function. This is in contrast with results in *Drosophila* and mammals, where some core clock genes exhibit very high amplitude oscillations (Hughes *et al*. 2009, Hughes *et al*. 2012, Li *et al*. 2015).

### Canonical clock genes and comparison with Drosophila

The canonical clock genes were examined for rhythmicity both at the transcript level and via an additional RAIN analysis at the gene level. The q-values (FDR adjusted p-values) for the canonical clock genes are shown in supplementary table S3. We found a rather limited evidence for rhythmicity in these genes which included pdp1e (q ~ 0.1, LL and DD), shaggy (q < 0.1, DD), and Clk (q ~ 0.1, LL). At a less stringent FDR (1 < 0.2), per, cyc, Dbt and cwo were rhythmic in DD, while cry and cyc, were oscillating in LL.

The most strongly associated cluster for the primary transcript (most highly expressed) of each gene is also shown in supplementary table S3, providing evidence that some clock genes are associated with clusters with rhythmic trends. For comparison between splice variants and conditions, median expression levels of the canonical clock genes and their transcripts for both DD and LL are shown in supplementary table S4.

We compared the transcripts identified as cycling in *Nasonia* heads with the transcripts identified as cycling in *Drosophila* heads. For these purposes, we used a list of genes identified in a meta-analysis study of *Drosophila* circadian microarray data as being rhythmically expressed in either LD or DD (Keegan *et al*. 2007). Of 173 genes identified as rhythmic in *Drosophila*, 33 genes (Supplementary table S5) were found to also be rhythmic in *Nasonia* (either in LL or DD, q < 0.1), no more than would be expected by chance (p = 0.11, hypergeometric test).

### Functions of rhythmic genes

To capture the general functions that rhythmic genes may fulfil in *Nasonia*, we tested a broader set of rhythmic genes (< 0.2 FDR in RAIN) for GO term overrepresentation (Davies and Tauber, 2015a), revealing 94 GO terms overrepresented for genes rhythmic in DD (including ‘response to light stimulus’, ‘proteasome complex’, and ‘generation of neurons’, Supplementary table S6) and 123 terms for genes rhythmic in LL (including ‘locomotion’,’proteasome complex’, and ‘response to external stimulus’, Supplementary table S7), 25 of which were shared between both conditions (Figure 3). Shared terms include terms related to neurons, signal transmission, and responses to stimuli. Notably, all four *Nasonia* opsins were found to exhibit similar transcriptional profiles in LL and DD, with low expression in the morning and high expression in the evening.

**Figure 3.**
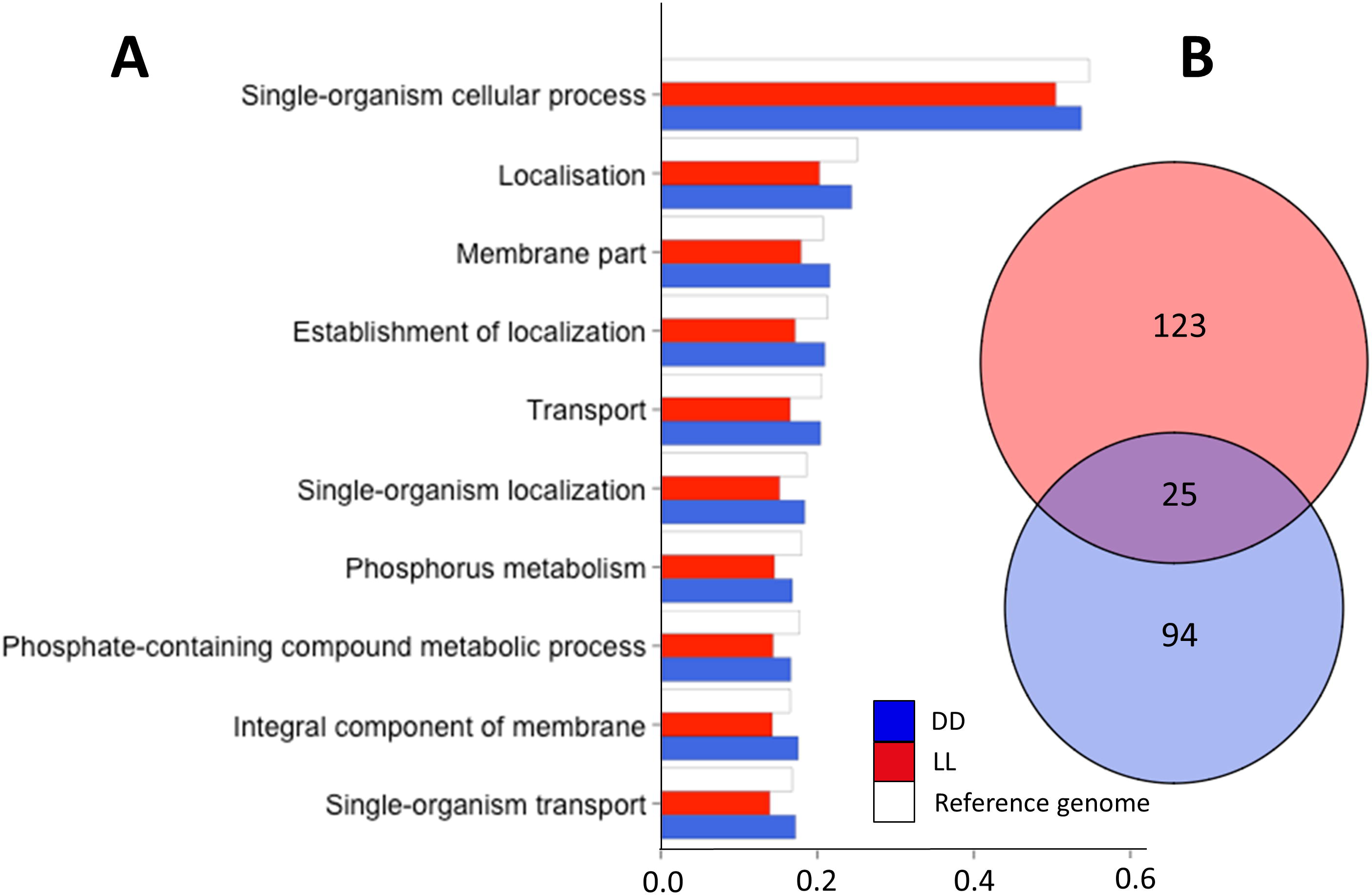
Enrichment of GO terms among cycling transcripts. (**A**) Bar plot of 10 top overrepresented GO terms (by gene proportion) for both DD and LL rhythmic genes. (**B**) Euler diagram showing the overlap of overrepresented terms in DD (blue) and LL (red).

It has previously been demonstrated that the timing of different (or indeed opposing) biological processes can be controlled through the circadian regulation of groups of genes (Sancar *et al*. 2015, Zhang *et al*. 2014). Unsupervised clustering methods have previously been established as a useful method for functional characterisation of circadian genes (Nguyen *et al*. 2014). To establish whether temporal separation of functions occurs in *Nasonia*, we therefore returned to the expression clustering analysis. Firstly, we employed hypergeometric tests to identify clusters with an overrepresentation of rhythmic genes (Figure 4, Supplementary table S8 and S9). Clusters which were found to have a significant rhythmic component (q < 0.05, supplementary tables S8 and S9) were analysed for overrepresented GO terms. Examples of clusters with enriched functions include clusters DD7 and LL20 which are significantly enriched for catalytic activity GO terms, especially genes involved in the proteasome, and clusters DD24 and LL6 which are both involved in circadian and neural processes. Other clusters (DD1 and DD2) did not turn up any overrepresented GO terms and are thus likely comprised of genes with a wide range of functions. Transcriptional differences between constant darkness and constant light To examine whether differences in circadian period seen in locomotor activity between DD and LL could also be detected in transcriptional rhythms, we fitted parametric models with a range of periods to transcripts rhythmic in both conditions (q < 0.1). For those transcripts with statistically significant fits to the model in both conditions (q < 0.1, see Methods), we took the period with the best fit and compared these periods between conditions. Overall, transcripts in LL showed a significantly (p < 3.9e-09, Wilcoxon rank sum test) shorter (median 24) period than those in DD (median 25.4), mirroring the behavioural differences in period.

**Figure 4.**
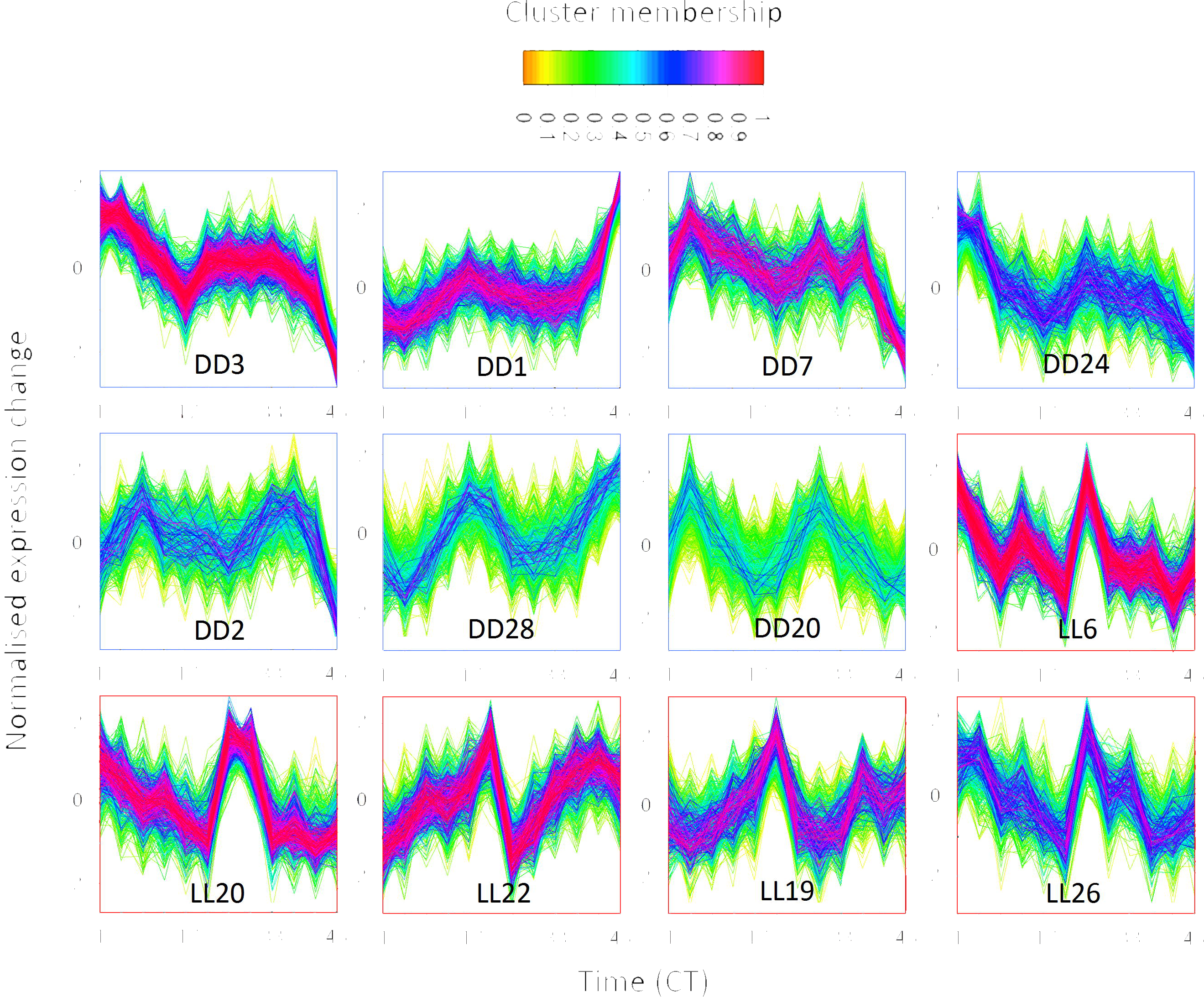
Normalised expression of clusters with significant (q < 0.01) overrepresentations of rhythmic genes. Each transcript profile in each cluster is coloured by that gene’s membership of the cluster.

We have also tested for differential expression between DD and LL. In the absence of biological replicates, we analysed differential expression using a fold change approach. We used 1.5 fold change as a cut-off for differential expression (Dalman *et al*. 2012), yielding 1,488 genes expressed higher in DD than LL and 971 genes expressed higher in LL than DD (Figure 5). Genes more highly expressed in DD were significantly enriched (q < 0.01) for genes involved in various forms of catalytic activity (Supplementary table S10), including the vast majority of proteasome genes (>75%). Genes more highly expressed in LL were enriched for a small number of terms including ‘plasmalemma’ and ‘sequence specific DNA binding’ (Supplementary table S11).

**Figure 5.**
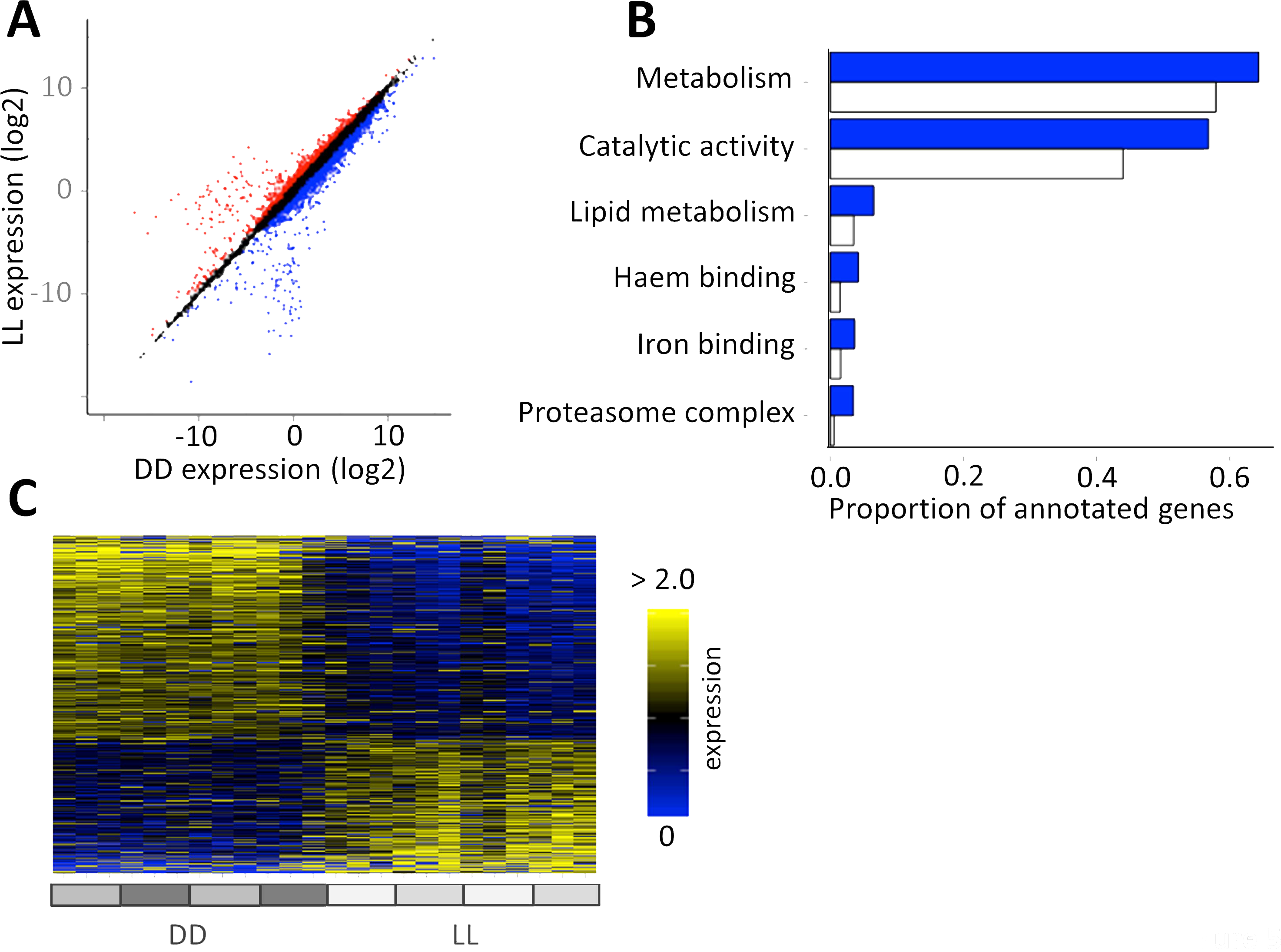
Comparison of the DD and LL transcriptomes. (**A**) FPKM (log2) expression of transcripts in DD (x axis) and LL (y axis), showing genes classified (> 1.5 median fold change) as differentially expressed up in DD (blue) and up in LL (red). (**B**) Selected overrepresented (q < 0.01) GO terms for genes more highly expressed in DD. (**C**) Heatmap showing median-normalised expression for differentially expressed transcripts, in DD (left) and in LL (right), sorted by fold change.

## Discussion

This study provides the first insights into global transcriptional oscillation in *Nasonia*. With RNA-seq, we profiled the circadian transcription of >26,000 transcripts in *Nasonia* in either DD or LL. At a relatively stringent FDR (q < 0.1), we identified 1,057 cycling transcripts in DD and 929 cycling transcripts in LL. These transcripts correspond to a cycling fraction of 6.7% and 5.9% of all transcripts analysed in DD and LL respectively. These figures are comparable to cycling fractions reported in various organisms and tissues, generally between 2% and 10% of the transcriptome (Michael and McClung, 2003).

In both conditions, cycling transcripts were found to cycle at low amplitudes (mostly < 2 fold) and with a limited, bimodal, range of phases. This is in contrast to microarray/RNA-seq studies in *Drosophila*, where transcripts were found to cycle with a broader range of phases (Rodriguez *et al*. 2013) and studies in both mammals and *Drosophila*, which have identified a group of high amplitude (> 4-fold) cycling genes among the low-amplitude majority (Akhtar *et al*. 2002). High amplitude cyclers typically include clock genes (Akhtar *et al*. 2002, Hughes *et al*. 2012). The low oscillations of the *Nasonia* head transcriptome render the expression profiles of the canonical clock genes difficult to resolve (Covington *et al*. 2008). This issue may also contribute to the discordance between the various circadian microarray studies in *Drosophila* (Keegan *et al*. 2007).

An emerging property of the circadian transcriptome in *Nasonia* is the temporal separation of function by phase (Fig 2). Notably, genes involved in catalytic activity were strongly overrepresented in morning-peaking transcripts. This is in line with other studies which show catalytic activity confined to the morning in fungi (Sancar *et al*. 2015), in agreement with a general observation that an important (or even primary) function of circadian clocks (Hurley *et al*. 2015) is to temporally separate catabolism and anabolism. Although we did not detect an overrepresentation of anabolic genes within the cyclic transcripts, expression clusters DD10 and LL24 (Supplementary figures S1 and S2) did show strong overrepresentation (Supplementary tables S12 and S13) for genes involved in cytosolic ribosomal genes (q < 3.e-56) and cellular anabolism (q < 2e− 06). These clusters exhibit an antagonistic expression pattern to the expression clusters containing the catabolic genes, suggesting that catabolism and anabolism are indeed separated by the circadian clock in *Nasonia*.

The comparison of expression between LL and DD reveals that a majority of genes involved in the proteasome and a broader set of genes involved in catabolism, are more highly expressed in DD than LL. As turnover rates of clock proteins have shown to be coupled with changes in the circadian period (Syed *et al*. 2011, He and Liu, 2005), up-regulation of the proteasome may provide an explanation for differences in period observed between DD and LL.

Although the similarity of genes which cycle in *Drosophila* and *Nasonia* is rather low, the functions fulfilled by CCGs in *Nasonia* are similar to the functions filled by CCGs in *Drosophila*. Examples of functions shared by CCGs in the *Drosophila* and *Nasonia* heads are: various aspects of metabolism (Rodriguez *et al*. 2013, Ueda *et al*. 2002, Ceriani *et al*. 2002, Claridge-Chang, *et al*. 2001), phototransduction (Ueda *et al*. 2002, Rodriguez *et al*. 2013), synaptic/nervous functions (McDonald and Rosbash, 2001b, Ceriani *et al*. 2002, Claridge-Chang *et al*. 2001), oxidoreductase activity (Claridge-Chang, *et al*. 2001), mating behaviour (Rodriguez, *et al*. 2013), and immunity (McDonald and Rosbash, 2001b, Ceriani, *et al*. 2002).

We identified cycling of genes involved in response to light, particularly all four *Nasonia* opsins. These opsins, along with associated gPCRs, cycle with a similar phase and are all more highly expressed in LL than in DD (Supplementary figure S6). Daily and circadian changes in opsin expression have been demonstrated in other organisms (e.g. mice (Bowes *et al*. 1988), zebrafish (Li *et al*. 2005), honeybee (Sasagawa *et al*. 2003)), and opsin expression is generally found to be up-regulated in response to light (Yan *et al*. 2014). Characterising the opsins in *Nasonia* is likely to provide insights into the light input pathway into the clock, particularly as *Nasonia* does not possess other obvious light input candidate genes such as *Drosophila*-like CRY1 (Bertossa *et al*. 2014) or Pteropsin (Velarde *et al*. 2005) (Supplementary figure S6).

### Data availability

We have made the expression profile for each transcript in both conditions available on WaspAtlas (Davies and Tauber, 2015b). Data have been archived in the NCBI short read archive (SRA), with accession number(s) [].

## Methods

### Maintenance and sample collection

Stocks of *Nasonia vitripennis* (strain AsymCX) were maintained at 25°C on blowfly pupal hosts in 12:12 light:dark cycles. To obtain male wasps for experiments, groups of eight females were isolated at the yellow pupal stage and transferred onto fresh hosts upon eclosion. The resulting male progeny were collected upon eclosion and moved onto vials with a 30% sucrose agar medium, in groups of 20. During entrainment (four full days in an LD 12:12 cycle) and collection, wasps were kept in four light boxes in the same incubator at 19°C. Starting at CT1, wasps were collected every four hours and snap-frozen in liquid nitrogen and immediately transferred to −80°C. Wasps were collected sequentially from light box to light box every four hours to minimise disturbance of wasps, and so that wasps were collected from each light box once every 16 hours, thereby minimising the effect of variations within light boxes. Temperature and light recordings were taken during the experiment, and can be viewed in Supplementary file S2. To verify that wasps entrained correctly to the experimental conditions and that free-running behaviour was as expected, individual male wasps were isolated and locomotor activity was monitored. Behavioural recordings of individual male wasps in experimental conditions can be seen in Supplementary figure S7, ruling out behavioural differences caused by inter light box variations in light intensity in LL, though not transcriptional differences.

### RNA extraction, sequencing, and read mapping

RNA was extracted from pooled groups of 50 heads for each sample, using Trizol RNA extraction protocol, and followed by clean-up using the RNAeasy spin column kit (Qiagen). Samples were polyA selected and sequenced at Glasgow Polyomics (University of Glasgow, United Kingdom) on the Illumina NextSeq500 platform, resulting in approximately 20 million 75bp paired-end reads per sample.

Read mapping was achieved with Tophat2 (v2.1.0)(Trapnell *et al*. 2012) against the *Nasonia* Nvit_2.1 NCBI annotation. As the purpose of this study was not to identify novel splice variants or improve on existing annotation, novel junction detection was disabled for accurate quantification of known transcripts. Mean mapping efficiency was above 90% for both conditions (Supplementary table S14). Read quantification was performing using the DEseq normalisation method (Anders and Huber, 2010). All 24 samples from both conditions were grouped together to allow comparison between as well as within conditions.

### Expression profile clustering

Isoform expression profiles were first filtered to include only those isoforms with no missing values at any time-point in either condition. Expression values were standardised using the ‘Standardise’ function in Mfuzz (Kumar and E Futschik, 2007). The ‘cselection’ function in Mfuzz was used to select an appropriate c-value for the c-means clustering (default parameters; m=1.25). Based on this analysis, thirty fuzzy clusters were generated for each condition using the fuzzification parameter m=1.25.

### Rhythmic expression analysis

RAIN (Thaben and Westermark, 2014) was used on all filtered isoforms (i.e. those with no missing values at any timepoint) in either condition to detect rhythmic isoforms at a period of 24 hours. As a non-parametric method, RAIN only facilitates detection of rhythmic isoforms with periods which are a multiple of the sample resolution (in this case 4 hr). The p-values produced by RAIN were corrected to q-values using the Benjamini-Hochberg method (Benjamini and Hochberg, 1995). This method was repeated using expression values for genes rather than transcripts for the clock gene analysis (i.e. the summed expression values for all known transcripts of a particular gene).

Maximum fold changes in expression were calculated by normalising percondition expression values by the median value and calculating the ratio from the lowest expression over 48 hours to the highest. Reliably quantified transcripts are defined as those those transcripts where the absolute FPKM value is 5 or above at all timepoints, the threshold for this set at a similar level to other analyses (Hughes *et al*. 2012).

To analyse the period of rhythmic transcripts, we fitted parametric waveforms with a variety of periods (20 to 28 hrs in steps of 0.2 hrs) to all transcripts identified as rhythmic (q < 0.1) in both conditions. This FDR threshold is in line with, or more strict, than thresholds chosen in other similar studies (Hughes *et al*. 2012, Huang *et al*. 2013, Keegan *et al*. 2007). Those transcripts (85 in total) which showed a significant (q < 0.1) fit to the model in both conditions were analysed in terms of their best fitting period.

GO term overrepresentation was performed in WaspAtlas (Davies and Tauber, 2015b) using the Nvit_2.1 NCBI annotation dataset. All hypergeometric tests were performed within R using the ‘phyper’ function. Clusters with rhythmic components were identified by collapsing the fuzzy clusters into hard clusters using the ‘cluster’ property of the Mfuzz object, performing hypergeometric tests to identify clusters with enrichment for rhythmic transcripts. Thirty tests were performed for each condition (i.e. for all clusters), and were corrected per-condition using the Benjamini-Hochberg method in R (R Development Core Team, 2008).

For comparison to microarray studies, orthologs for *Drosophila melanogaster* were obtained from a meta-study of circadian microarray data (Keegan *et al*. 2007). The 214 obtained FlyBase identifiers were converted to the latest identifiers using the validation tool, resulting in 218 unique identifiers (the increase in identifiers can be attributed to previous identifiers referring to multiple genes in the current annotation). Orthologs for these *Drosophila* genes were obtained through WaspAtlas, retrieving orthologs for 135 genes which mapped to 173 unique *Nasonia* genes due to gene duplications, etc. This set of 173 genes was compared with the number of genes with rhythmic transcripts that would be expected by chance using a hypergeometric test.

### Phylogenetic analysis of opsin genes

Opsin genes were searched for using NCBI BLASTP using six species; *Apis mellifera, Bombyx mori, Drosophila melanogaster, Mus musculus, Nasonia vitripennis, and Homo sapiens, using the Nasonia Lop1* protein sequence as a query. BLAST results were inspected and 7e-19 was chosen as an appropriate cut-off to include all opsin sequences. Sequences were aligned by ClustalW in MEGA (Tamura *et al*. 2007) and a maximum likelihood tree generated using default parameters. Duplicated sequences were manually removed, and sequences renamed for display on the tree. Full protein name to shortened display name translations can be found in supplementary table S15.

